# Cosmid based mutagenesis causes genetic instability in *Streptomyces coelicolor*, as shown by targeting of the lipoprotein signal peptidase gene

**DOI:** 10.1101/049320

**Authors:** John T Munnoch, David A. Widdick, Govind Chandra, Iain C. Sutcliffe, Tracy Palmer, Matthew I Hutchings

## Abstract

Bacterial lipoproteins are a class of extracellular proteins tethered to cell membranes by covalently attached lipids. Deleting the lipoprotein signal peptidase (*lsp*) gene in *Streptomyces coelicolor* results in growth and developmental defects that cannot be restored by reintroducing the *lsp*. We report resequencing of the genomes of the wild-type M145 and the *cis*-complemented Δ*lsp* mutant (BJT1004), mapping and identifying secondary mutations, including an insertion into a novel putative small RNA, *scr6809*. Disruption of *scr6809* led to a range of developmental phenotypes. However, these secondary mutations do not increase the efficiency of disrupting *lsp* suggesting they are not *lsp* specific suppressors. Instead we suggest that these were induced by introducing the cosmid St4A10Δ*lsp* as part of the Redirect mutagenesis protocol, which transiently duplicates a number of important cell division genes. Disruption of *lsp* using no gene duplication resulted in the previously observed phenotype. We conclude that *lsp* is not essential in *S. coelicolor* but loss of *lsp* does lead to developmental defects due to the loss of lipoproteins from the cell. Significantly, our results indicate the use of cosmid libraries for the genetic manipulation of bacteria can lead to unexpected phenotypes not necessarily linked to the gene or pathway of interest.

## Introduction

Bacterial lipoproteins are essential for building and maintaining the cell envelope and also provide a key interface with the external environment ^1–3^ Most lipoprotein precursors are exported as unfolded polypeptides via the Sec (general <u>sec</u>retory) pathway but others can be exported via the twin arginine transport (Tat) pathway, which is typically utilised for the transport of fully folded proteins ^4–6^ The signal peptides of lipoproteins closely resemble other types of bacterial Sec and Tat signal peptide but they contain a characteristic lipobox motif, typically L_−3_-A/S_−2_-G/A_−1_-C_+1_, relative to the signal cleavage site, in which the cysteine residue is essential and invariant. The lipobox motif allows putative lipoproteins to be easily identified in bacterial genome sequences^3,7^.

Following translocation, lipoprotein precursors are firstly modified by covalent attachment of a diacylglycerol molecule, derived from a membrane phospholipid, to the thiol of the conserved lipobox cysteine residue via a thioether linkage. This reaction is catalysed by an enzyme named Lgt (<u>L</u>ipoprotein diacyl<u>g</u>lycerol <u>t</u>ransferase) and results in a diacylated lipoprotein. Lsp (<u>L</u>ipoprotein <u>s</u>ignal <u>p</u>eptidase) then cleaves the signal sequence immediately upstream of the lipidated cysteine to leave it at the +1 position. These early steps in lipoprotein biogenesis are highly conserved and unique to bacteria making them potential targets for antibacterial drug development ^2,8^. In Gram-negative bacteria and Gram-positive Actinobacteria, lipoproteins can be further modified by addition of an amide-linked fatty acid to the amino group of the diacylated cysteine residue at the mature N-terminus. This final step is catalysed by the enzyme Lnt (<u>L</u>ipoprotein <u>n</u>-acyl<u>t</u>ransferase) and results in triacylated lipoproteins. In Gram-negative proteobacteria, Lnt modification is a pre-requisite for the recognition of lipoproteins by the Lol machinery, which transports lipoproteins to the outer membrane ^2,9^ but its function in monoderm Gram-positive bacteria is not known. Members of the Gram-positive phyla Firmicutes and Mollicutes also N-acylate lipoproteins despite lacking Lnt homologues and *S. aureus* can diacylate or triacylate individual lipoproteins in an environmentally dependent manner ^10–14^. These studies suggest that triacylation of lipoproteins in Gram-positive bacteria has an important role in their natural environment but is dispensable *in vitro*. Loss of Lnt activity in *Streptomyces* bacteria has no obvious effect on fitness or lipoprotein localisation *in vitro* but it does have a moderate effect on virulence in the plant pathogen *Streptomyces scabies*, supporting the idea that it has environmental importance ^15^.

We previously characterised all four steps of the lipoprotein biogenesis pathway in *Streptomyces* spp. (Figure 1) ^5,15^, which is one of the best studied genera in the Gram-positive phylum Actinobacteria. Our key findings are (i) that Tat exports ~20% of lipoprotein precursors in streptomycetes; (ii) they N-acylate lipoproteins using two non-essential Lnt enzymes; (iii) *Streptomyces coelicolor* encodes two functional copies of Lgt which cannot be removed in the same strain; (iv) *lsp* mutants can be isolated at low frequencies but they acquire spontaneous secondary mutations which might be *lsp* suppressors. It was recently reported that Lgt is essential in *Mycobacterium tuberculosis*, which is also a member of the phylum Actinobacteria, and that *lgt* deletion in the fast-growing species *Mycobacterium smegmatis* is accompanied by spontaneous secondary mutations ^16^. Natural product antibiotics that target the lipoprotein biogenesis pathway include globomycin, made by *Streptomyces globisporus* ^2^ and antibiotic TA made by *Myxococcus xanthus* ^1,16^. Both inhibit Lsp activity and are lethal to *Escherichia coli* but TA resistance arises through spontaneous *IS3* insertion into the *lpp* gene, which encodes an abundant lipoprotein that attaches the *E. coli* outer membrane to the peptidoglycan cell wall ^16,17^. Over-expressing *lsp* also confers TA resistance in both *E. coli* and *M. xanthus*, and the latter encodes additional Lsp homologues within the TA biosynthetic gene cluster ^17^.

**Figure 1.**
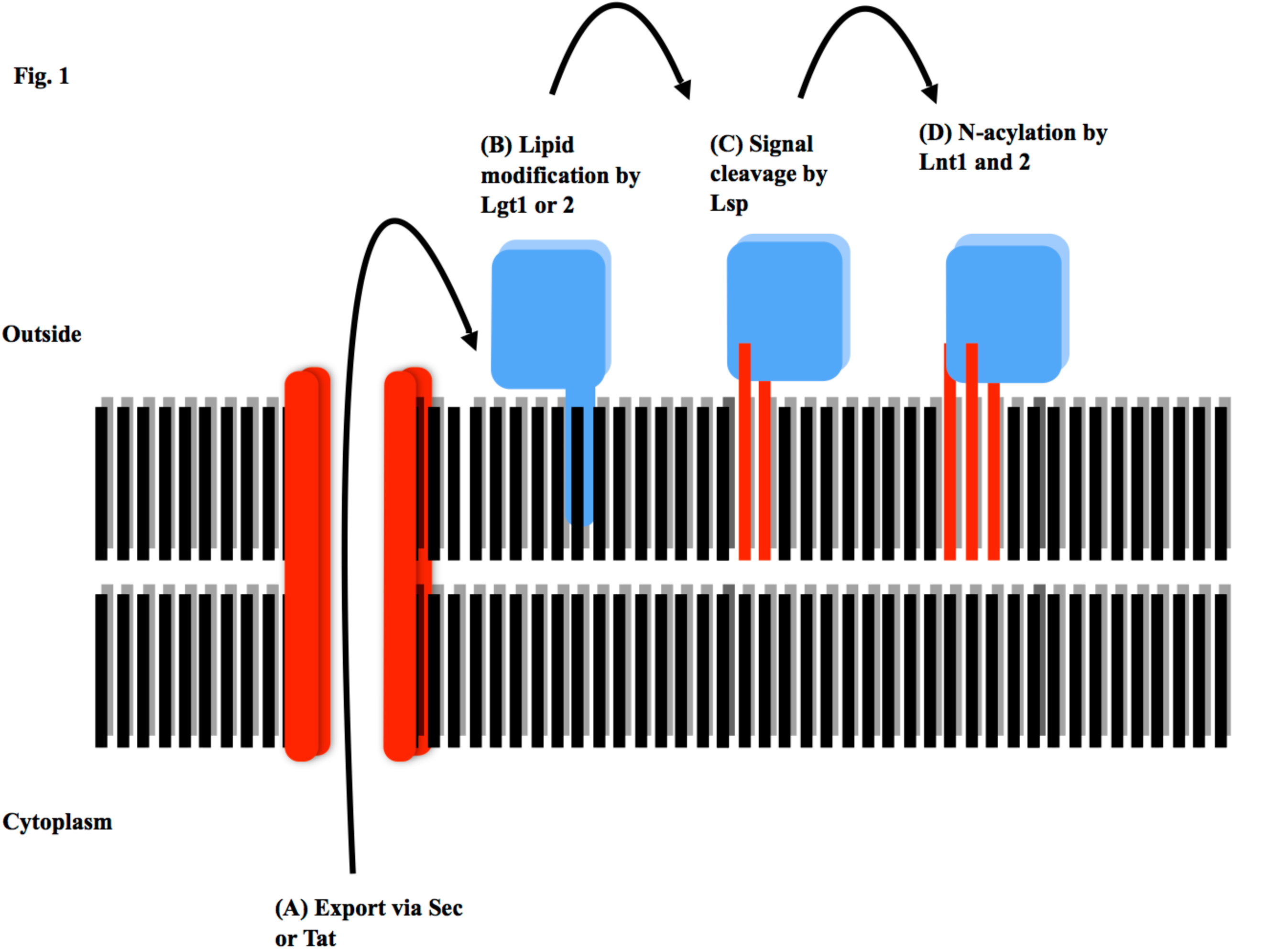
Lipoprotein biogenesis in *Streptomyces coelicolor*. Approximately 80% of precursor lipoproteins in *S. coelicolor* are translocated via the general secretory (Sec) pathway with around 20% being translocated by the twin arginine transport (Tat) pathway (A). Following translocation across the cytoplasmic membrane they are diacylated on the thiol of the lipobox (+1) cysteine residue by Lgt1 or Lgt (B) and then the signal sequence is cleaved by Lsp immediately upstream of that modified cysteine (C). Lnt1 then adds a third acyl chain to the amino group on the +1 cysteine to produce a triacylated lipoprotein (D). Lnt2 is not essential for triacylation *in vitro* but appears to increase its efficiency. The function of the N-acyl modification is not yet known.

Deletion of *S. coelicolor lsp* results in very small and flat colonies that are delayed in sporulation and these *lsp* mutants could not be fully complemented even by reintroducing the *lsp* gene to its native locus. Although both *cis* and *in trans* complementation restored lipoprotein biogenesis and sporulation it did not restore the wild-type growth rate ^5^. There are two likely reasons for this: either *lsp* is essential and the mutant strains acquire secondary suppressor mutations, or the Redirect PCR targeting method that we used to delete the *lsp* gene resulted in chromosomal rearrangements and mutations independent of *lsp*. Here we provide evidence to support the second hypothesis and we demonstrate that introduction of the cosmid carrying an ~40 kb region of the *S. coelicolor* chromosome, including *lsp*, from *E. coli* to *S. coelicolor* transiently duplicates cell division and cell wall biosynthesis genes which leads to secondary mutations including disruption of a putative small RNA. We further confirm that *lsp* is non-essential but deletion of the *lsp* gene does lead to growth and developmental delays and the over-production of the antibiotic actinorhodin in *S. coelicolor*, as observed previously. These phenotypes must therefore be due to the loss of lipoproteins from its cytoplasmic membrane.

## Results

### Mapping secondary mutations in the *cis* complemented *Δlsp* strain BJT1004

We previously reported that the *S. coelicolor* Δ*lsp* mutant BJT1001 cannot be complemented even by restoring *lsp* to its native locus ^5^. Since *cis* complementation should effectively restore the genome to wild-type this suggests that other spontaneous mutations have occurred during the genetic manipulations. To test this we Illumina sequenced the genomes of the parent strain *S. coelicolor* M145 and the *cis-*complemented Δ*lsp* strain BJT1004 using two independent companies (GATC Biotech and The Genome Analysis Centre). Across the four sequence samples a total of 51 unique single nucleotide polymorphisms (SNPs) were detected (Supplementary Table S1 online) as well as a chromosomal rearrangement in BJT1004 that is not present in the parent strain M145 (Figure 2(A-B) and Supplementary Text S1 and S2 online). Of the 51 SNPS, 13 are unique to one of the BJT1004 sequences, with 4 residing inside coding regions. However of all of these only one SNP occurs in both BJT1004 sequences and this is in the intergenic region between *sco5331* and *sco5332*. In the single chromosomal rearrangement, the *IS21* insertion element (genes *sco6393* and *sco6394*) has inserted into the intergenic region between the *sco6808* and *sco6809* genes and this was confirmed by PCR (Figure 2(A-C)). Although this might affect the downstream promoter of *sco6808*, which encodes a putative regulator, deletion of *sco6808* (using vector pJM010) had no effect on growth or development under standard laboratory conditions (Figure 3(A)). The intergenic position of *IS21* in BJT1004 suggested it might disrupt a non-coding RNA and examination of RNA sequence data for *S. coelicolor* M145 confirmed the presence of a 189 nt transcript initiating 107 bp upstream of *sco6808* and reading into the last 82 nucleotides of the *sco6809* gene (data from GSM1121652 and GSM1121655 RNA sequencing; Supplementary Text S1 and 2 online; Figure 2(A-B)). Following convention we named this putative small RNA *scr6809* for *<u>S</u>. <u>c</u>oelicolor* <u>R</u>NA 6809. Deletion of the *scr6809* (pJM012) sequence (without disrupting either the *sco6808* or *sco6809* coding sequences) resulted in a range of phenotypes from colonies that look like wild-type to non-sporulating bald and white mutants defective in aerial hyphae formation and sporulation, respectively, antibiotic overproducers and small slow growing colonies. Restreaking the Δ*scr6809* colonies (double crossovers) with wild-type appearance again gave rise within the next generation to a range of colonies with different morphologies, including growth and developmental defects (Figure 3(B)). Colonies with mutated morphologies would retain that morphology in subsequent generations indicating another situation where spontaneous secondary mutations are arising. A previous report showed that a *sco6808* deletion mutant had accelerated production of actinorhodin and undecylprodigiosin as well as precocious spore formation on R5 medium ^18^. There was no observable difference between the wild-type and Δ*sco6808* strains under the growth conditions used here but disruption of *sco6808* in strain BJT1004 resulted in an improvement in sporulation (Figure 3(A)). We suspect this difference is based on the recovery of the *scr6809* loci to wildtype as result of the double crossovers between the chromosome of BJT1004 and the Δ*sco6808* deletion cosmid St1A2Δ*sco6808*. This was also seen for St1A2Δ*sco6811* disruptions in each background (not shown).

**Figure 2.**
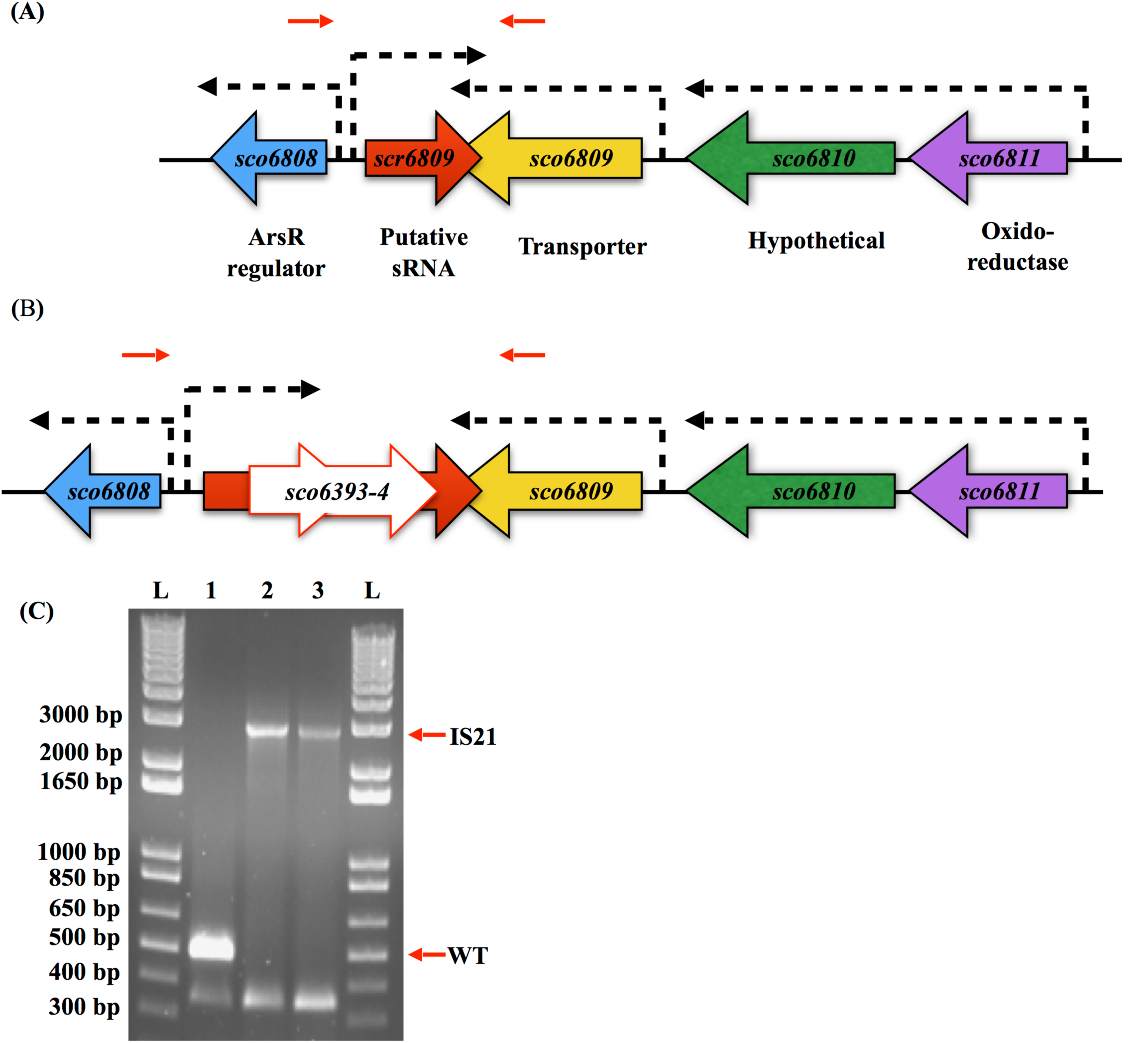
*IS21* insertion into *scr6809*. The *sco6811-08* region of the *S. coelicolor* M145 genome contains 4 genes (*sco6811* (Purple), *sco6810* (green), *sco6809* (yellow) and *sco6809* (blue) and a putative sRNA *scr6809* (large red) along with 3 putative promoters (broken arrows). Representations of the WT loci (A) and that sequenced from BJT1004 (B) indicate where an IS21 element (*sco6393* and sco6394) has inserted within *scr6809*. PCR verification of the IS21 insertion with primers JM0093 and JM0094 (small red arrows) was carried out (C) using M145 (lane 1), BJ1001 (lane 2) and BJT1004 (lane 3) genomic DNA. Lanes marked L contain the size ladders (Invitrogen 1kb plus DNA ladder), lane 1 contains the PCR product using wild-type M145 DNA (514 bp), lane 2 contains the PCR product using Δ*lsp* strain BJT1001 DNA and lane 3 contains the PCR product using genomic DNA from the cis complemented Δ*lsp* strain BJT1004 (both 2884 bp.

**Figure 3.**
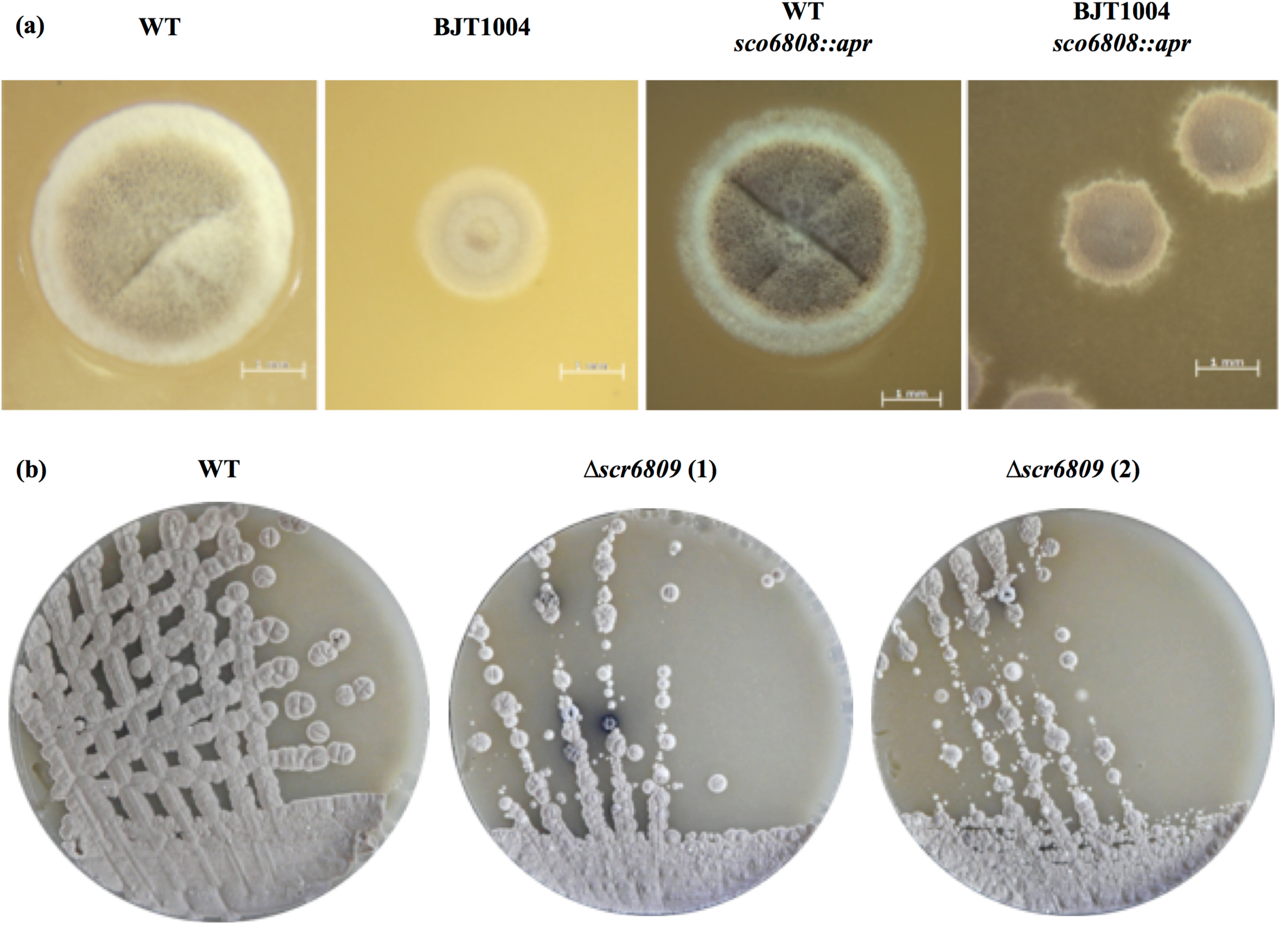
Analysis of the *IS21* disrupted genomic region in *S. coelicolor*. Colony morphology (A) shows that deletion of *sco6808* has no obvious effect on growth or development in wild-type M145 but does partially restore sporulation in BJT1004 (recovery of *scr6809*). Disruption of *scr6809* in M145 results in a range of pleiotropic morphological and developmental phenotypes (B).

To determine whether *IS21* insertion into *scr6809* is induced by deletion of *lsp*, we isolated ten more non-clonal *lsp* mutants by introducing cosmid St4A10Δ*lsp* (pJM014) into wild-type strain M145 and then PCR amplified the intergenic region between *sco6808* and *sco6809*. The size of the PCR products matched the predicted wild-type size and indicated that none of these *lsp* mutants contain an *IS21* insertion suggesting that the original observation is not specific to *lsp* mutants (Figure 4). Consistent with this conclusion, the frequency with which *lsp* mutants could be isolated was not increased in BJT1004 relative to M145 suggesting that none of the mapped mutations in BJT1004 suppress fitness defects that arise from deleting Δ*lsp*. Attempts at over-expressing *scr6809* using pJM017 in *S. coelicolor* M145, *S. scabies* 87-22 and *S. venezualae* ATCC 10712 resulted in no observable phenotype but as the same vectors failed to prevent accumulation of developmental phenotypes in the Δ*scr6809* strain, this suggests that functional *scr6809* is not expressed from these vectors.

**Figure 4.**
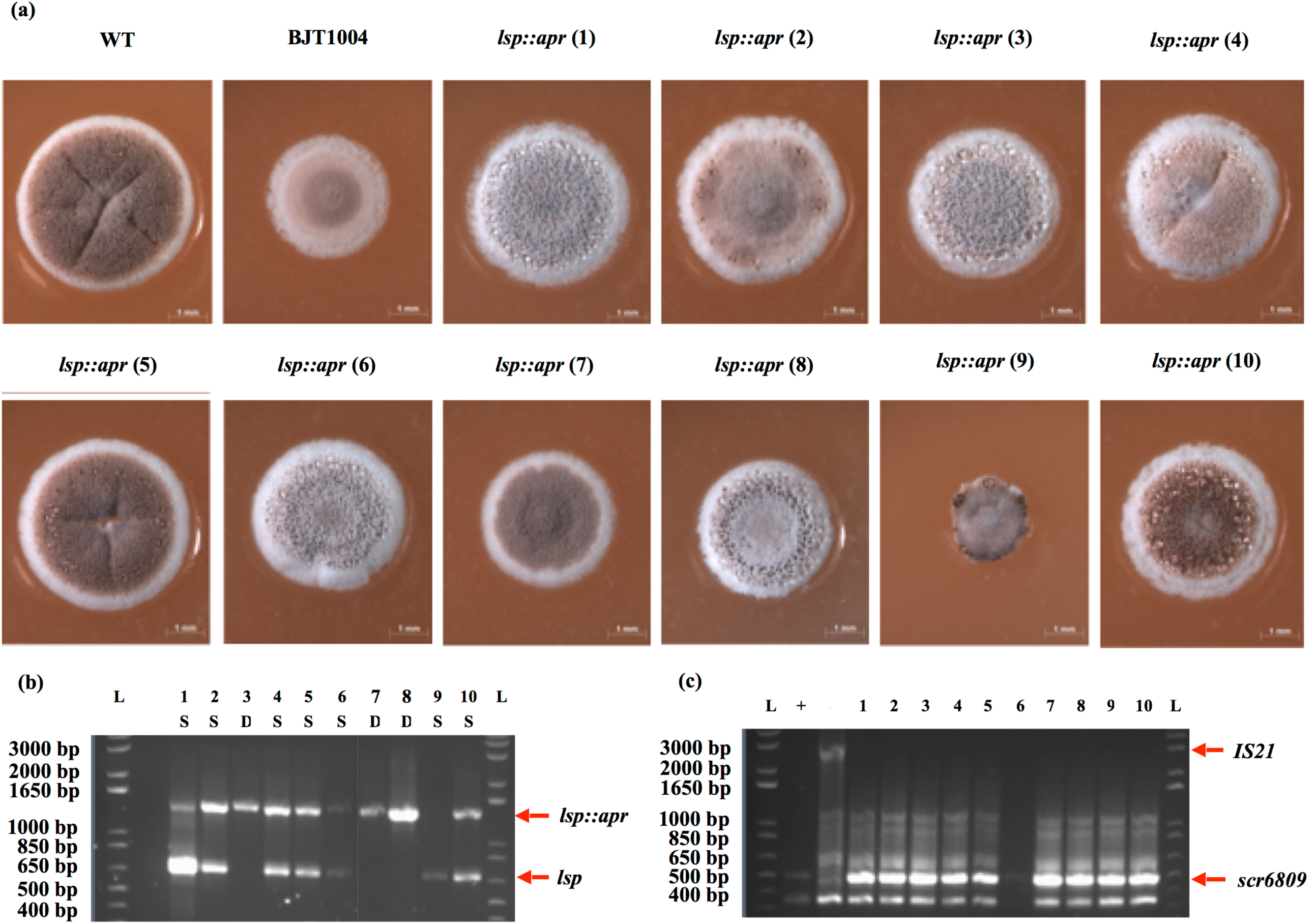
New *lsp* mutants generated using Redirect do not contain the *IS21* mutation. Colony morphology of mutants *lsp::apr* 1-10 (corresponding to strains JTM008.01 – JTM008.10), both single (n=7, colonies 1-2, 4-7 and 9-10) and double crossovers (n=3, colonies: 3 and 7-8) show a range of phenotypes (A). PCR of the *lsp* loci indicates colonies are either a single (WT and/or mutant band) or double (mutant band only) crossovers (WT = 687 bp, mutant = 1447 bp).

Cumulatively these results suggested that deletion of *lsp* does not result in secondary mutations and prompted us to hypothesise that these accumulate as a result of duplicating cell division genes on cosmid St4A10 which was used to delete *lsp*. These results further suggest a role for *scr6809* in *S. coelicolor* differentiation, although there is no obvious link to *lsp* and so this was not pursued further here.

### Introduction of wild-type St4A10 results in a pleiotropic phenotype

The Redirect PCR-targeting method uses *E. coli* as a host strain for an *S. coelicolor* cosmid library which can be used to make targeted deletions ^19,20^. The Redirect method was used to PCR-target the *lsp* gene *sco2074* on cosmid St4A10, which contains a ~40 kb region of the *S. coelicolor* genome spanning genes *sco2069-2104* (Supplementary Table S2 online). Conjugation of St4A10Δ*lsp* into *S. coelicolor* transiently duplicates all the genes on that cosmid (except *lsp*) and because this region includes many important cell division genes (*ftsZ*, *ftsQ*, *ftsW*, *ftsI* and *ftsL*) and essential cell wall synthesis genes (*murG*, *murD*, *murX*, *murF* and *murE*) we reasoned that over-expression of these genes, rather than deletion of *lsp*, is responsible for the spontaneous secondary mutations and the resulting pleiotropic phenotype. To test this idea we introduced an origin of transfer into the wild-type St4A10 cosmid backbone and then conjugated this cosmid into strain M145 and selected for single cross-over events where the whole cosmid is integrated into the chromosome, thus duplicating the *S. coelicolor* genes on St4A10. Analysis of these single crossover strains, maintained on kanamycin to select for the cosmid, revealed them to be genetically unstable, with many initially appearing similar to the observed Δ*lsp* phenotype, i.e. small and delayed in sporulation (Figure 5). However, they lack the characteristic Δ*lsp* overproduction of the blue antibiotic actinorhodin and colonies from this M145::St4A10 strain also acquired more significant developmental issues upon prolonged maintenance and restreaking on MS agar containing kanamycin (not shown). This suggests they accumulate spontaneous secondary mutations as a direct result of carrying St4A10 and that the observed Δ*lsp* phenotype is at least in part due to duplication of the genes on cosmid St4A10. This is consistent with the fact that complementation of Δ*lsp* restored lipoprotein biogenesis but did not restore wild-type colony morphology ^5^.

**Figure 5.**
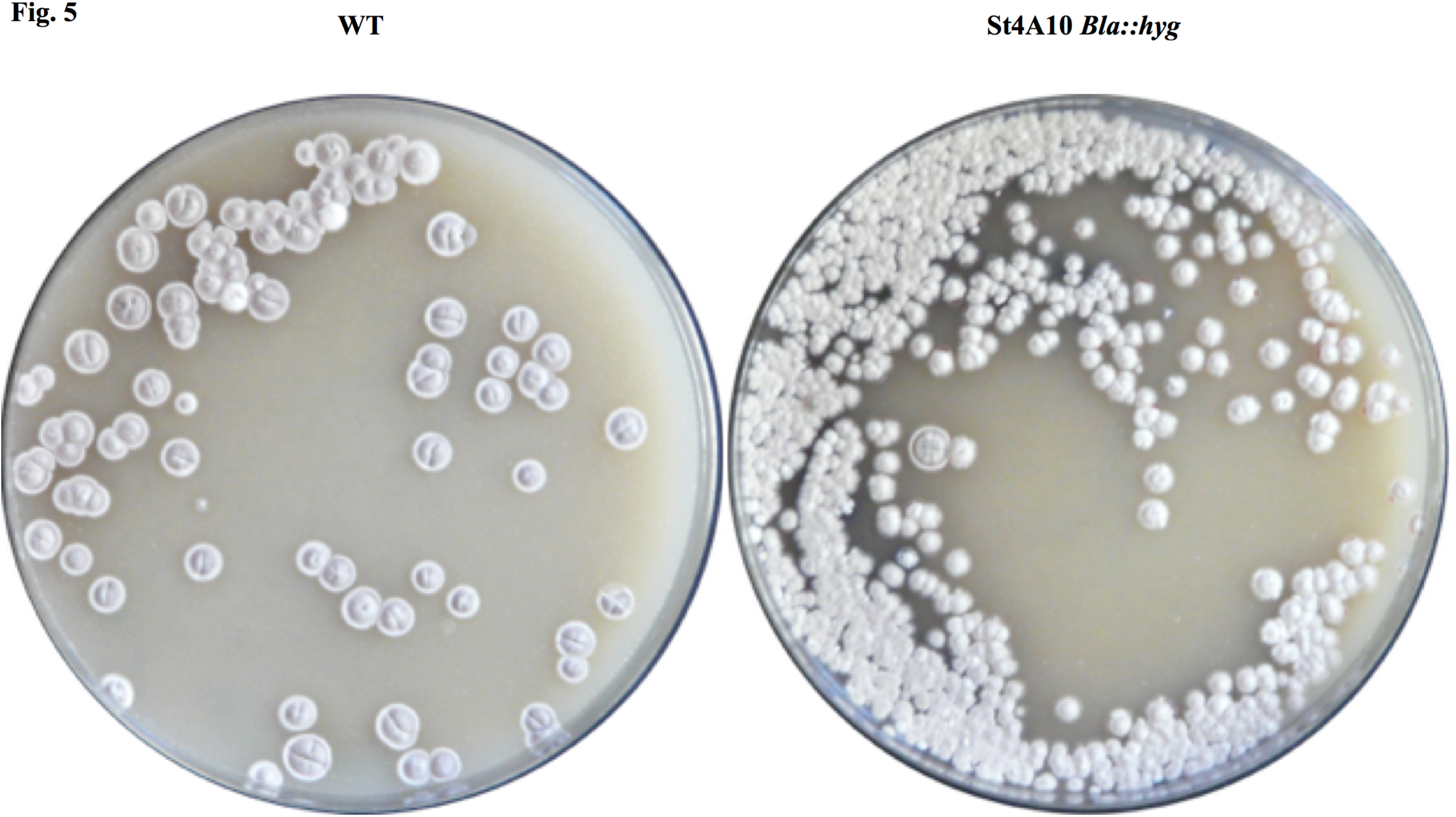
Introduction of wild-type St4A10 causes a pleiotropic phenotype. Conjugation of M145 with St4A10 *bla::hyg* results in non-wildtype phenotypes similar to those observed in the St4A10 *lsp::apr* single crossovers.

### Targeted deletion of *lsp* results in a small colony phenotype

To test how much deletion of *lsp* contributed to the phenotype of BJT1001 (the Δ*lsp* strain generated using Redirect) we undertook a targeted disruption of *lsp* in wild-type strain M145 using a suicide vector, which does not duplicate or affect any other coding sequences. The *lsp* suicide vector, pJM016 (Table 1), was introduced into wild-type *S. coelicolor* by conjugation and ex-conjugants were selected by growing on MS agar plates containing apramycin. Following introduction of the pJM016, two colony types were observed (Figure 6), one with wild-type appearance while the others were small colonies that over-produce actinorhodin, reminiscent of the *lsp* mutant BJT1001. PCR testing of the genomic DNA of both morphotypes revealed that those with the wild-type colony morphology have a wild-type fully functioning *lsp* gene whereas those with a small colony phenotype have disruptions in *lsp* caused by pJM016. PCR amplification followed by sequencing of the loci in the small colony variant revealed an interesting and unexpected recombination event had occurred: the vector and almost all the *lsp* gene have been removed such that all that remains is the apramycin resistance cassette (Supplementary Text S4 and S5 online). These data confirm that *lsp* is not essential in *S. coelicolor* but loss of Lsp does result in a growth and developmental defect and overproduction of the blue antibiotic actinorhodin as observed previously ^5^.

**Figure 6.**
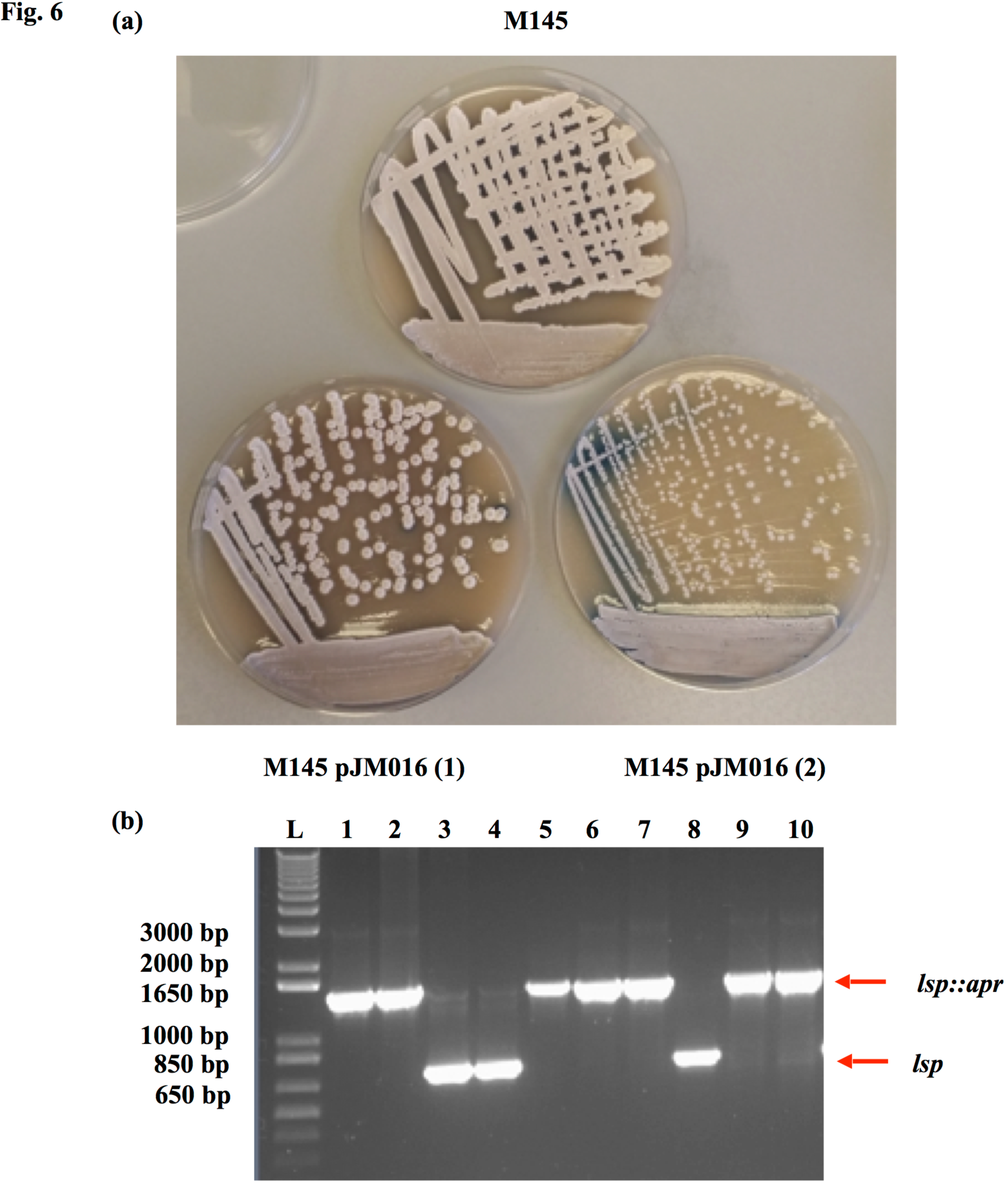
Targeted disruption of *lsp* using a suicide vector results in a small colony phenotype that overproduces actinorhodin. To test how much of the BJT1001 phenotype is due to loss of *lsp* we disrupted the *lsp* gene using a suicide vector which does not affect or duplicate any other target genes. Plate images (A) show two distinct phenotypes following insertion of the suicide vector (pJM016) into M145, either a wildtype appearance (M145 pJM016 (1), n=3 corresponding to JTM018.03-04 and 08) or a small colony phenotype over producing actinorhodin (M145 pJM016 (2), n=7, corresponding to strains JTM018.01-2, 05-07 and 09-10) similar to our original observation of the *lsp* phenotype ^5^.

**Table 1.**
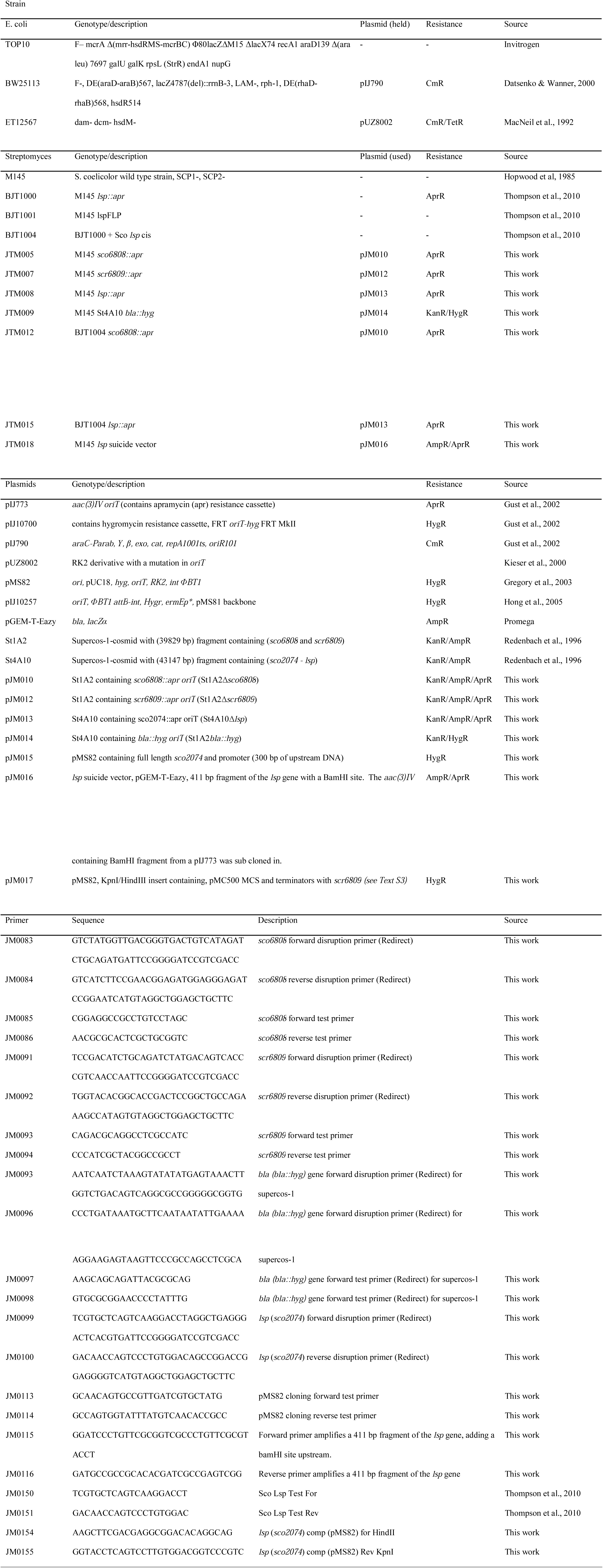
**Strains, plasmids and primers**.

## Discussion

The pleiotropic nature of the original *S. coelicolor* Δ*lsp* strain BJT1001 resulted primarily from the introduction of cosmid St4A10, most likely caused by the over-expression of cell division and cell wall biosynthesis genes carried on that cosmid (Supplementary Table S2 online). It seems likely, but is not proven, that this led to the secondary mutations we observed in this strain. These secondary mutations do not make it easier to delete *lsp* suggesting they are not *lsp-*specific suppressors. Genetic manipulation has always been challenging in *Streptomyces* bacteria and the Redirect PCR targeting method has been a significant development but this work should be a cautionary tale to others to consider the effects of using large insert cosmid libraries in the genetic manipulation of bacteria. Recent advances in CRISPR/Cas9 editing of *Streptomyces* genomes ^21^ negate the need for a cosmid library and these techniques will accelerate research into the basic biology of *Streptomyces* and other filamentous actinomycetes. This is vital because the secondary metabolites derived from these bacteria still represent a major underutilised reservoir from which new antibiotics and other bioactive natural products can be discovered. Moreover, the identification here of the novel small RNA *scr6809* and demonstration that its deletion results in a range of growth and developmental defects add to the growing appreciation ^22–24^ of the significance of small RNAs in streptomycetes.

## Materials and Methods

### Bacterial strains and culture conditions

All primers, plasmids and strains used are listed in Table 1. Strains were routinely grown as previously described ^5^ following the recipes of Kieser et al., (2000). *E. coli* was grown in LB or LB–NaCl for Hygromycin selection and *S. coelicolor* M145 and its derivatives were grown on Soya Flour Mannitol (SFM) medium to study growth and development or LB culture for genomic isolations.

### Gene deletions and complementation

Gene deletions were carried out following the Redirect method of PCR-targeting ^26^ as previously described Hutchings *et al*. (2006). Disruption of *lsp (sco2074::apr)* on cosmid St4A10 (pJM013, St4A10Δ*lsp*) using the pIJ773 apramycin disruption-cassette and *sco6808 (sco6808::*a*pr)* and *scr6809 (scr6809::apr)* on cosmid St1A2 (pJM010 - St1A2Δ*sco6808* and pJM012 - St1A2Δ*sco6808* respectively) using primers JM0101-2, JM0083-84 and JM0091-2 respectively were confirmed by PCR using primers JM0150-1, JM0085-6 and JM0093-4 respectively. Introduction of the wild-type cosmid St4A10 was facilitated by introducing an *oriT* by disruption and replacement of the Supercos-1 backbone *bla* resistance gene (pJM014 – St4A10*bla::hyg)* using primers JM0095-6 and the hygromycin disruption cassette from pIJ10701, confirmed using primers JM0099-100. The *lsp* suicide vector pJM016 was produced by introducing a 411 bp fragment of the *lsp* gene with an N-terminal *Bam*HI site, amplified with primers JM0117-8 and cloned into pGEM T-Eazy. The *Bam*HI site was then used to subclone the *Bam*HI fragment from a pIJ773 digest, containing an *apr* disruption cassette. An overexpression construct, pJM017 was synthesised by Genscript to include the pMC500 MCS and terminators ^28^ with *scr6809* (sequence included in Supplementary Text S5 online). All constructs were subsequently conjugated into *S. coelicolor* following the method described by Gust et al. (2002).

### Genomic DNA isolation

Genomic DNA was isolated from M145 and BJT1004 following the Pospiech and Neumann (1995) salting out method as described by Keiser et al. (2000). Mycelium from a 30 ml culture was resuspended in 5 ml SET buffer containing 1 mg/ml lysozyme and incubated at 37°C 30-60 min. To this lysate, 140 μl of proteinase K solution (20 mg ml^−1^) was added, mixed, then 600 μl of 10% SDS added, mixed and incubated at 55°C for 2 h, with occasional mixing throughout. After this incubation 2 ml of 5 M NaCl was added, mixed and left to cool to 37°C before adding 5 ml chloroform, mixed at 20°C for 30 min. Samples were centrifuged at 4500 x g for 15 min at 20°C. The supernatant was removed to a fresh tube and DNA precipitated by adding 0.6 volumes of 100% isopropanol. Tubes were mixed by inversion and after at least 3 min DNA spooled out using a sterile Pasteur pipette. The DNA was rinced in 70% ethanol, air dried and dissolved in 1-2 ml TE buffer (10 mM Tris-HCl pH 7.8, 1 mM EDTA) at 55°C.

### Genome resequencing and secondary mutation identification

The isolated DNA from our wild-type *S. coelicolor* M145 parent strain and BJT1004 were sent to both GATC Biotech and The Genome Analysis Centre (TGAC) for 35bp paired end HiSeq Illumina sequencing. Assembly mapping and SNP identification was carried out with MIRA (Chevreux et al., 2004) using the reference genome NC_003888 (Bentley et al., 2002) as a scaffold for mapping each of the resequenced genomes. Putative SNPs were detected in each sample independently reporting the SNP position, the nucleotide change, the number of reads that sequence the region, those containing wild-type or mutated nucleotides and a percentage change. Each set of results was then compared by eye to determine the likely hood that a SNP was real by number of reads containing the mutation and its presence in each sample. Larger mutations (rearrangements) were identified in the same fashion.

### Microscopy

Brightfield images were acquired using a Zeiss M2 Bio Quad SV11 stereomicroscope. Samples were illuminated from above using a halogen lamp images captured with an AxioCam HRc CCD camera. The AxioVision software (Carl Zeiss, Welwyn Garden City, UK) was used for image capture and processing.

## Acknowledgments

We thank Marie Elliot for sharing unpublished RNA sequence data for *Streptomyces coelicolor* M145. This work was supported by a NERC PhD studentship to JTM and BBSRC grants BB/F009429/1 and BB/F009224/1 to MIH and TP, respectively.

### Author contribution

JTM designed and carried out the experiments, JTM ICS TP MIH designed experiments, JTM DAW ICS TP MIH analysed data and wrote the manuscript, DAW prepared DNA for sequencing, GC analysed sequencing data.

### Additional information

**Supplementary information** accompanies this paper at:

**Competing financial interests:** The authors declare no competing financial interests.

